# Genome sequencing of the vermicompost strain *Stenotrophomonas maltophilia* UENF-4GII and population structure analysis of the *S. maltophilia* Sm3 genogroup

**DOI:** 10.1101/2021.08.12.455975

**Authors:** Francisnei Pedrosa-Silva, Filipe P. Matteoli, Hemanoel Passarelli-Araujo, Fabio L. Olivares, Thiago M. Venancio

## Abstract

The *Stenotrophomonas maltophilia* complex (Smc) is a cosmopolitan bacterial group that has been proposed an emergent multidrug-resistant pathogen. Taxonomic studies support the genomic heterogeneity of Smc, which comprises genogroups exhibiting a range of phenotypically distinct strains from different sources. Here, we report the genome sequencing and in-depth analysis of *S. maltophilia* UENF-4GII, isolated from vermicompost. This genome harbors a unique region encoding a penicillin-binding protein (*pbp*X) that was carried by a transposon, as well as horizontally-transferred genomic islands involved in anti-phage defense via DNA modification, and pili glycosylation. We also analyzed all available Smc genomes to investigate genes associated with resistance and virulence, niche occupation, and population structure. *S. maltophilia* UENF-4GII belongs to genogroup 3 (Sm3), which comprises three phylogenetic clusters (PC). Pan-GWAS analysis uncovered 471 environment-associated and 791 PC-associated genes, including antimicrobial resistance (e.g. *bla*L1 and *bla*R1) and virulence determinants (e.g. *tre*S and *kat*G) that provide insights on the resistance and virulence potential of Sm3 strains. Together, the results presented here provide the grounds for more detailed clinical and ecological investigations of *S. maltophilia*.

## INTRODUCTION

Vermicomposting is a non-thermophilic biodegradation technique used to manage organic waste [1]. The process involves synergistic interactions between earthworms and microorganisms to biodegrade different organic waste into a humus-like material known as vermicompost, a nutrient-rich organic amendment used to enhance soil microbial diversity and plant development [2, 3]. Vermicomposts harbor soil bacteria from various genera, such as *Bacillus, Pseudomonas, Serratia,* and *Stenotrophomonas*, which may engage in beneficial interactions with plants [4, 5].

*Stenotrophomonas maltophilia* is a Gram-negative bacillus found in a wide range of natural habitats, including water sources, soils, rhizospheres, animal microbiotas, including humans [6, 7]. *S. maltophilia* has also been used as part of bioremediation and biocontrol strategies [6, 8]. However, *S. maltophilia* has been reported as a global multidrug resistant opportunistic pathogen associated with significant mortality rates of up to 37.5%, mainly because of bacteremia and respiratory tract infections in severely debilitated, immunosuppressed or chronic lung disease patients [9, 10]. *S. maltophilia* is intrinsically resistant to multiple classes of antibiotics, such as aminoglycosides, carbapenems, and macrolides, posing a therapeutic challenge and delay the administration of proper antibiotics [7].

*S. maltophilia* present high intraspecific variability [11] and, along with the closely related species *S. pavanii*, comprise the *S. maltophilia* complex (Smc) [12–14]. Phylogenic studies based on multilocus sequencing typing (MLST) and whole-genome sequencing revealed the organization of Smc members in several genogroups [15–17]. A study using whole-genome multilocus sequence typing (wgMLST) and average nucleotide identity (ANI) analyses showed the genetic organization of Smc in 23 monophyletic genogroups with different virulence and resistance characteristics [18]. Among those, genogroup 3 (Sm3) exhibits a myriad of phenotypically distinct strains from different sources. These strains remain poorly explored, hampering investigations on the genetic determinants underlying the physiology and niche occupation of *S. maltophilia* isolates.

Here we present the genome sequencing of *S. maltophilia* UENF-4GII, the first *S. maltophilia* isolated from vermicompost. *S. maltophilia* UENF-4GII belongs to Sm3 and harbors a set of interesting horizontally-transferred regions, including genomic islands (GIs) involved in phage resistance via DNA modification, and pili glycosylation. ANI analysis and SNP-based phylogenetic reconstructions allowed us to reclassify six publicly available *Stenotrophomonas* spp. genomes as *S. maltophilia* from Sm3. We also used the *S. maltophilia* UENF-4GII genome with those of other 66 *S. maltophilia* isolates to thoroughly characterize of the population structure, virulence, and resistance profiles of Sm3. Finally, a pangenome-wide association study (pan-GWAS) of Sm3 allowed us to identify genes associated with niche occupation and phylogenetic clusters.

## METHODS

### Bacterial isolation and identification

The bacterium was isolated from mature manure vermicompost produced from cattle manure at Universidade Estadual do Norte Fluminense Darcy Ribeiro, Brazil. In summary, serial dilutions (10^−1^ to 10^−7^) were performed on a solution prepared by adding 10 g of vermicompost in 90 mL of saline (8.5 g·L^−1^ NaCl), followed by shaking for 60 min. Then, 100 μL of the final dilution (10^−7^) were taken and spread on plates containing solid Nutrient Broth (NB) with 8 g·L^−1^ of NB and 15 g·L^−1^ of agar in 1 L of distilled water. After incubation at 30 °C for 7 days, different colony types could be identified and, for purification, individual colonies were transferred to Petri plates with Dygs solid media acquired from Vetec (São Paulo, Brazil). After isolation and purification on Dygs solid medium, a yellowish, circular, convex elevation, punctiform and smooth surface bacterial colony was selected. Light microscopy revealed a Gram-negative strain and the presence of rod-shaped motile was confirmed under phase contrast microscopy. This isolate, named UENF-4GII, was stored in a 16 mL glass flask containing 5 mL of Nutrient Broth solid medium covered with mineral oil and later grown in liquid Dygs medium under rotatory shaker at 150 rpm and 30 °C for 36 h.

### Genome sequencing and annotation

Genomic DNA was extracted using QIAamp^®^ DNA Mini Kit (Qiagen) and quantified with an Agilent Bioanalyzer 2100 instrument (Agilent, California, USA). Paired-end libraries were previously prepared with the TruSeq Nano DNA LT Library Prep (Illumina) and sequenced on an Illumina HiSeq 2500 sequencing system at the Life Sciences Core Facility (LaCTAD; UNICAMP, Campinas, Brazil). Sequencing reads (2 x 100 bp) were assembled *de novo* with SPAdes v.10.3.1 [19] and scaffolded with Gfinisher v.1.4 [20], using as references the complete genome of *S. maltophilia* JV3 (GCF_000223885.1) and an alternative assembly generated with Velvet 1.2.10 [21]. The assembly statistics were assessed with QUAST v.3.0 [22]. Genome completeness was assessed with BUSCO v.4.0 [23], using the *Gammaproteobacteria* dataset as reference. PlasmidSpades [24] was used to predict plasmids. The assembled genome was annotated with the NCBI Prokaryotic Genome Annotation Pipeline (PGAP) [25]. The UENF-4GII genome was deposited into DDBJ/EMBL/GenBank under the accession number JABUNQ000000000. Genes involved in antimicrobial metabolite biosynthesis were predicted using antiSMASH [26]. Insertion sequences (ISs) and GIs were predicted with ISEscan v.1.5.4 [27] and Islandviewer4 [28], respectively. Bacteriophage signatures were analyzed with PHASTER [29].

### Genome similarity assessment

We downloaded 641 genomes of *Stenotrophomonas* available in the RefSeq database in January, 2021 [30, 31]. Genome completeness was assessed with BUSCO 4.0 [23], using quality score ≥ 90% and the *Gammaproteobacteria* dataset as reference. We excluded assemblies with more than 500 contigs. A preliminary genome distance estimation analysis of isolate UENF-4GII and RefSeq genomes was performed using Mash [32] and a distance tree was generated using MASHtree v.0.50 [33]. All-against-all ANI based on MUMmer alignment (ANIm) was performed with pyani v.0.27 [34]. To assess the concordance between ANI and Mash estimates, we performed a linear correlation analysis using Pearson’s correlation coefficient. For this analysis, the Mash distances were converted into Mash scores (1-Mash distance) to allow a direct comparison with ANI values.

### Pangenome analysis

The Sm3 genogroup pangenome was performed with Roary v.3.6, using 95% identity threshold to determine orthologous [35]. Core genes were aligned with MAFFT v.7.394 [36]. SNPs were extracted from the core-genome alignment using SNP-sites v.2.3.3 [37] and maximum likelihood phylogenetic reconstructions were performed with IQ-tree [38], with ascertainment bias correction under the model GTR+ASC. The bootstrap support was evaluated using the ultrafast bootstrap method with 1000 replicates [39]. The resulting phylogenetic tree was visualized with iTOL v4 [40].

As a complementary approach, we performed a pangenome-wide association study (pan-GWAS))on the on the Sm3 dataset using Scoary [92]. The pan-GWAS was computed using the Roary output to find genes associated with isolation source (clinical or non-clinical) and to establish which genes were typical of each phylogenetic cluster (PC), while correcting for population structure using the core-genome phylogenetic tree (command –n tree). False-discovery rate was estimated by Benjamini–Hochberg adjusted p-value provided in Scoary. We only reported the results with specificity > 70% and Benjamini–Hochberg corrected p-value < 0.05. The binary heatmaps of trait-associated genes were rendered using R package tidyverse [41].

### Virulome and resistome analysis

Virulence and antimicrobial resistance genes were predicted using Usearch v.11.0.667 screened against the Virulence Factors of Pathogenic Bacteria Database (VFDB) and the Comprehensive Antibiotic Resistance Database (CARD). Minimum identity and coverage thresholds of 60% and 50% were used in these searches, respectively. The presence/absence profiles of virulence and resistance-associated genes were rendered using R package tidyverse [41].

## RESULTS

### Genome analysis of *S. maltophilia* UENF-4GII

During the characterization of culturable bacteria from mature cattle vermicompost, we identified a bacterium that was preliminarily characterized as *Stenotrophomonas* sp. using 16S rRNA sequencing. We submitted the genome of this isolate to whole-genome sequencing (see methods for details). The 30,093,445 paired-end reads and were assembled into 3 scaffolds, with an N_50_ of 1.3 Mbp, encompassing a total of 4.4 Mbp with 66.55% GC content. No plasmids were detected. BUSCO assessment recovered 437 (96.68%) single copy genes of *Gammaprotebacteria* dataset, supporting the completeness and high quality of the assembled genome. The genome harbors 3,932 protein-coding genes, 70 tRNA genes, and 2 rRNA operons.

The *S. maltophilia* UENF-4GII genome carries 18 complete IS elements: nine of the IS481 family, seven of the IS3 family and a single copy of the IS5 and IS21 families (Supplementary figure S1). In addition, we also found an incomplete prophage region that is closely related with a temperate bacteriophage from *Burkholderia pseudomallei* (Burkho_phi1026b) [42]. AntiSMASH prediction revealed that UENF-4GII possesses four gene clusters involved in the production of two bacteriocin-like compounds, an aryl polyene related to Xanthomonadin [43], and a non-ribosomal peptide synthetase (NRPS) gene cluster involved in the biosynthesis of siderophore (Supplementary figure S1). Siderophores are the most important iron uptake systems in *S. maltophilia* [44]. In NRPS gene cluster, we identified six genes (HRE58_11480 to 11500, and HRE58_11505) encoding an specific siderophore of *S. maltophilia* [45]. These enzymes are similar to those involved in enterobactin biosynthesis in enteric bacteria (e.g. *E. coli*) [44].

### Genome similarity of UENF-4GII

In order to investigate the genomic relatedness of UENF-4GII strain, we downloaded all available *Stenotrophomonas* genomes in Refseq database (n=641, January 2021), out of which 627 were retained after BUSCO quality filtering (see method for details). Genome identity was computed using Mash and ANI. The mash-distance phylogeny highlights the diversity of the *Stenotrophomonas* genus, corroborating the *S. maltophilia* complex organization in different genogroups (Figure 1A), as previously described by Gröschel et al. [18]. We identified 17 genogroups according to the distance estimation of well-characterized representative genomes, including the affiliation of UENF-4GII to Sm3, along with other 67 clinical and non-clinical strains (Supplementary table S1). In addition, this analysis allowed us to reclassify six *Stenotrophomonas* spp. as *S. maltophilia* Sm3 isolates, including the environmental strains *Stenotrophomonas* sp. DDT-1 (GCF_001580555.1) and *Stenotrophomonas* sp. Pemsol (GCF_003586545.1), reported as specific biodegraders of organochlorine pesticide (DDT) and polycyclic aromatic hydrocarbons (PAH), respectively. Mash results are supported by ANI analysis, which showed genomic identity above 95% within the same genogroup (Figure 1B). Moreover, the strong positive linear correlation (r = 0.98, p-value < 2.2e-16) between ANI and Mash results (Figure 1C) confirm a similar level of resolution for species delineation within the *Stenotrophomonas* genus.

**Figure 1.**
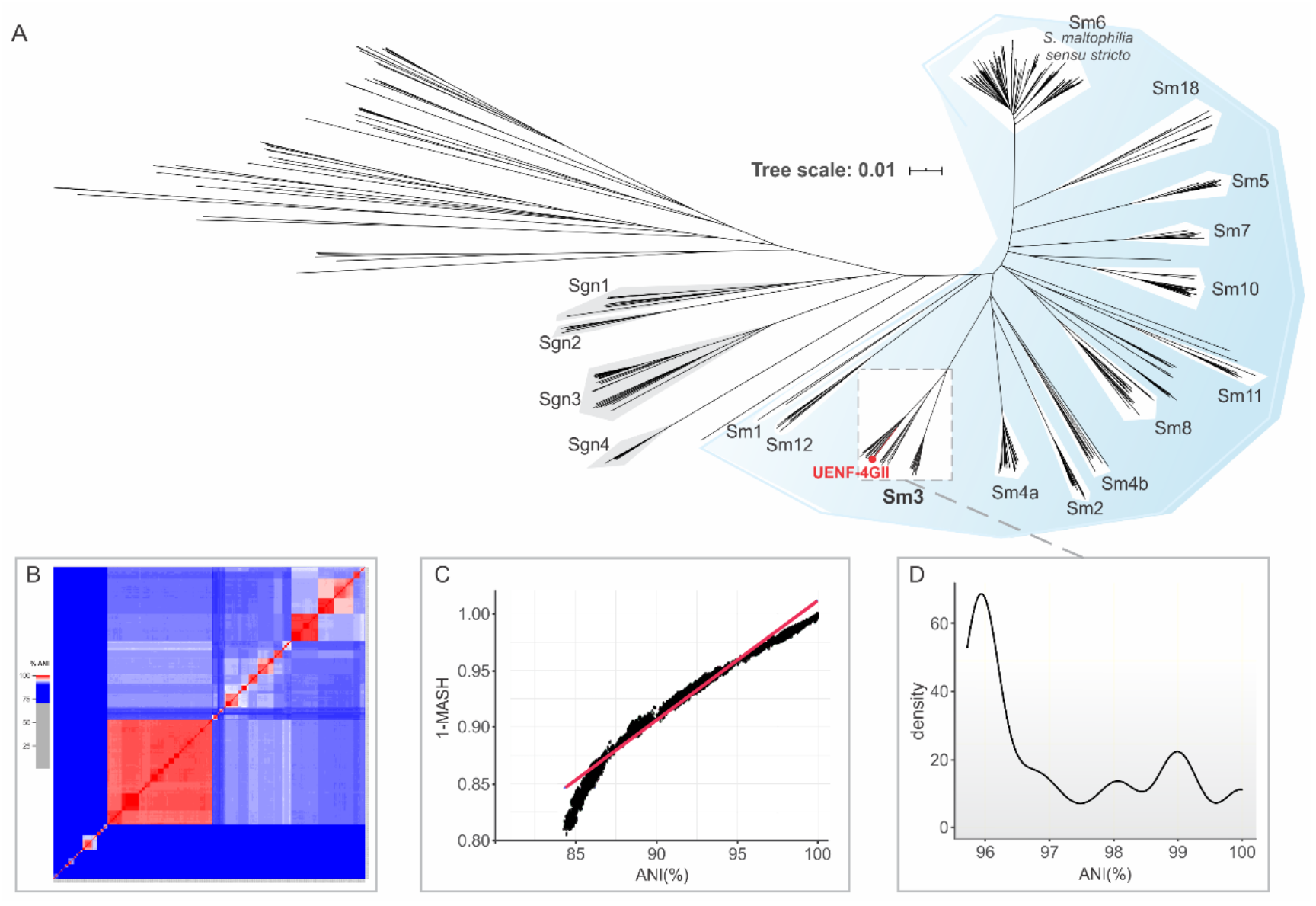
Genomic diversity of *Stenotrophomonas* genus. (A) Mash-distance-based phylogeny of *Stenotrophomonas*, built using 627 publicly genomes and that of UENF-4GII (red color). The *S. maltophilia sensu lato* clade is shaded in light blue. (B) Pairwise average nucleotide identity (ANI) calculated with 627 *Stenotrophomonas* genomes. Colors depict the degree of genome identity. (C) Correlation between ANI and Mash methods. (D) Density plot of pairwise ANI within the Sm3 genogroup.

### Pangenome analysis

The pangenome analysis of the 67 Sm3 isolates comprises 13,380 genes, with 2,754 core genes (i.e. present in at least 95% of the strains), 1,958 shell genes (present in 15% to 95% of the strains), and 8,668 cloud genes (present in up to 15% of the strains). The heap law estimate supports an open pangenome (alpha= 0.65) (Figure 2A), which typically reflects a high genetic diversity through the acquisition of exogenous DNA [46]. The genomic fluidity (ϕ) Sm3 was estimated in 0.18 (± 0.05), indicating that an average of 18% of unique gene families between in a given pair of genomes.

**Figure 2.**
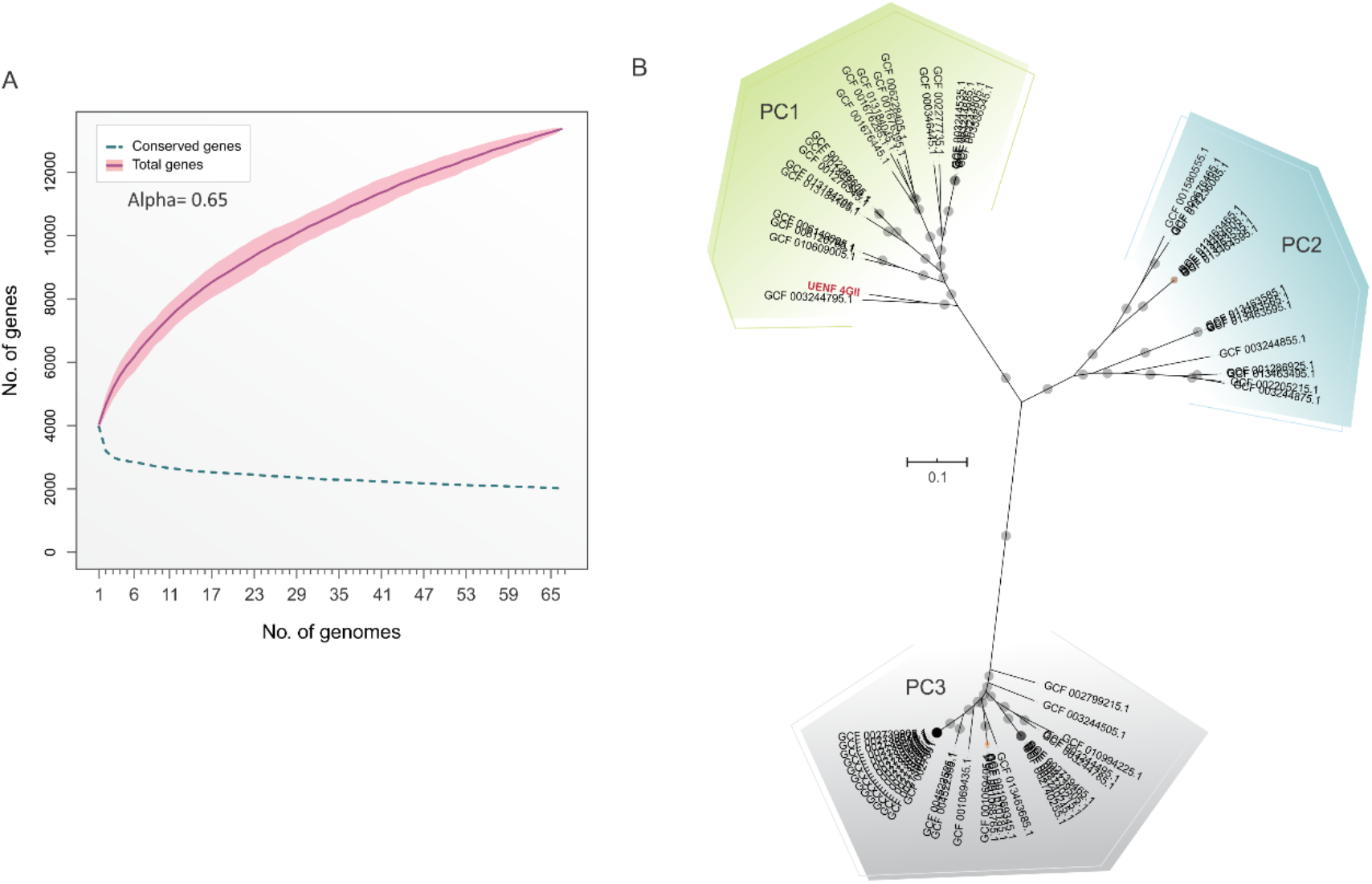
Pangenome and phylogeny of Sm3 genogroup. (A) Number of gene families in the Sm3 pangenome. The cumulative curve (in dark-red) and alpha value of the Heap law less than one (0.65) supports an open pangenome. (B) cgMLST of Sm3 genogroup showing three phylogenetic groups. SNPs extracted from the core genome were used to build a maximum likelihood phylogenetic tree using IQ-tree (see methods for details). Bootstrap values below and above 70% are represented by orange and gray points, respectively.

We also performed a maximum-likelihood phylogenetic reconstruction using SNPs extracted from core genes (i.e. core-genome multilocus sequencing typing, cgMLST). This analysis revealed that Sm3 comprises three distinct and highly supported phylogenetic clusters (PC) (Figure 2B). PC1 (n= 21) and PC2 (n= 15) encompass clinical and non-clinical strains, while PC3 (n= 31) has only clinical strains. Interestingly, 10 of the 21 PC1 strains are non-clinical, including *S. maltophilia* UENF-4GII.

We identified 75 unique genes in *S. maltophilia* UENF-4GII (Supplementary table S2), including *pbp*X (HRE58_12720), which encodes a low molecular weight penicillin-binding protein (LMW-PBP). This gene is closely located to other unique genes: a gene with unknown function (HRE58_12725) and two transposase genes (HRE58_12730, HRE58_12735), flanked by a perfect 26-bp inverted repeat (IR) sequence, supporting their acquisition by horizontal gene transfer (Supplementary figure 2). LMW-PBPs are important enzymes involved in cell-wall recycling and considered the major molecular targets for β-lactam antibiotics [47]. Alterations in the structure of LMW-PBPs are associated with reduced susceptibility to penicillin and other β-lactams and can induce the expression of β-lactamases [47–49]. The presence of an LMW-PBP composite transposon likely confers a competitive advantage for *S. maltophilia* UENF-4GII to thrive in microbial communities containing penicillin.

### UENF-4GII genomic islands

To further understand genome plasticity, we identified and analyzed 17 putative GIs (GI1-17) in *S. maltophilia* UENF-4GII, which probably represent recent horizontal gene transfers (Supplementary table S3). In addition, we compared these GIs with other Sm3 genomes (Figure 3). Among these regions, GI3 harbors an operon (HRE58_02625-02660) containing genes encoding GalE, HldD, and glycosyltransferases involved in lipopolysaccharide biosynthesis. This operon is present in 68% of the PC3 genomes. GI16 contains a class C beta-lactamase and GI10 have blue-light sensor genes that are absent in PC3 genomes.

**Figure 3.**
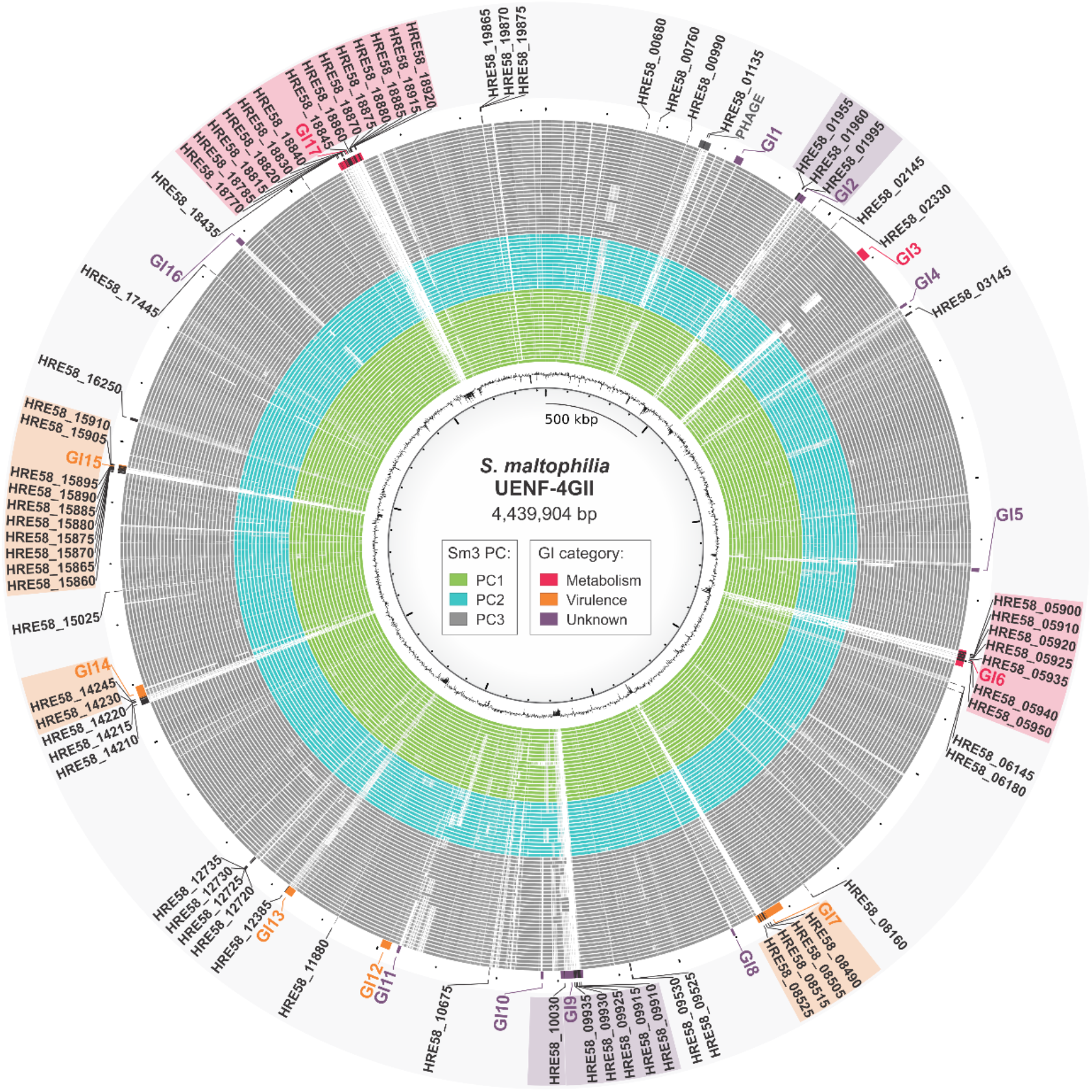
Circular genome representation of 75 unique genes and 17 genomic islands (GIs) identified in *S. maltophilia* UENF-4GII in comparison to other 66 Sm3 strains. The inner ring represents the UENF-4GII genome GC content. Green, blue and gray rings represent genomes from PC1, PC2 and PC3, respectively. The GIs were predicted and classified as metabolic (red), virulence (orange) or unknown (purple). Unique genes predicted within GIs were highlighted according to the GI category.

Interestingly, 60% of the *S. maltophilia* UENF-4GII unique genes are located within 7 GIs related with DNA metabolism (GI6 and GI17), virulence (GI7 and GI15) and unknown functions (GI2, GI9 and GI14). GI6 has 9 unique genes, including a restriction-modification (R-M) system with two type I restriction endonucleases (HRE58_05925 and HRE58_05935) and a DNA methyltransferase (HRE58_05920). R-M systems defend the bacterial genomes against bacteriophages and other types of exogenous genetic elements [50] and, hence, their acquisition might be adaptive by boosting resistance against phages [51, 52]. The GI17 has 41,433 bp and comprises 14 unique genes, including a DNA polymerase-like protein (HRE58_18840) and an operon containing an 8-oxoguanine DNA glycosylase (HRE58_18870), a nitroreductase (HRE58_18875), a 7-cyano-7-deazaguanine synthase (HRE58_18880; PreQ(0)), and a nucleoside 2-deoxyribosyltransferase (HRE58_18885). These genes are likely part of a repair system for oxidative stress-mediated DNA damage [53]. PreQ(0) might be involved in the insertion of 7-deazaguanine derivatives in the DNA [54], as part of a defense system against foreign DNA and phages [55].

Finally, 9 of 10 genes from GI15 are unique (Supplementary table S3). This island is involved in the biosynthesis and glycosylation of type IV pilin-like proteins. Hence, this island is likely an important virulence factor that mediate bacterial adherence to biotic and abiotic surfaces. Although pilin glycosylation has been associated with immune evasion of pathogenic bacteria [56], a recent study showed its role in the defense against phage infection [57]. Collectively, these GIs indicate that *S. maltophilia* UENF-4GII is equipped with different horizontally-acquired genes that increased its tolerance to foreign DNA.

### Pan-GWAS of Sm3 genogroup

We performed a pan-GWAS to identify genes associated with environment type (clinical and non-clinical; Supplementary table S1) and PCs in Sm3 using Scoary. We found 299 and 172 gene clusters associated with clinical and non-clinical environments, respectively (Supplementary figure S3, Supplementary table S4). Interestingly, 58% of the 471 environment-associated genes encode hypothetical proteins with unknown function.

Among the genes associated with clinical environments, we identified genes encoding a chemotaxis response regulator protein (*che*B), blue-light sensing protein (*blu*F), and trehalose sintase *tre*S, a crucial enzyme involved in trehalose biosynthesis. Several bacteria use trehalose as carbon source and stress protectant [58]. Trehalose metabolism has also been associated with the emergence of virulent human pathogens [59, 60]. Interestingly, *tre*S was also reported as a significant gene associated with clinical strains of the *S. maltophilia* complex [61]. Moreover, we identified significant associations with genes involved in bacterial resistance, including multidrug efflux pumps (*mdt*A and *mdt*C), macrolide efflux (*mac*A), aminoglycoside-modifying enzyme (*aac*), β-lactamases (*bla*L1 and *bla*L2), and a β-lactam sensor gene (*bla*R1). BlaL1 and BlaL2 are well known metallo-β-lactamases in *Stenotrophomonas*. They hydrolyze almost all β-lactams, are resistant to all clinically available β-lactamase inhibitors [62, 63] and have been show to increase pathogenicity in clinical settings [63, 64]. Among the genes associated with non-clinical environments, we also found genes involved in bacterial resistance, particularly in macrolide efflux (*mac*A and *mac*B), as well as three gene clusters involved in heavy metal resistance including copper (*cop*B), cobalt/zinc/cadmium (*czc*B), and mercury (*mer*R1).

As most non-clinical strains belong to PC1 and PC3 comprised only clinical strains, most of the environment-related genes are also linked with their respective phylogenetic groups. A total of 252, 97 and 442 gene clusters were associated, respectively, with PC1, PC2 and PC3 (Figure 4, Supplementary table S5). The majority (60%) of the PC-associated genes encoded proteins with unknown functions. PC1 was strongly associated (100% of sensitivity and specificity) with 13 gene clusters, including a type II toxin-antitoxin gene (*pas*I) and c-AMP phosphodiesterase encoding gene (*cpd*A) involved in stress resistance [65, 66]. Further, 11 gene clusters were strongly associated to PC2, including *bes*A, involved in iron metabolism [67], and *kat*G, related with oxidative stress resistance [68]. The PC3 presented 107 associated gene clusters, with 100% sensitivity and specificity. Among these, we identified genes involved in transcriptional regulation (*cue*R, *com*R, *rst*A, and *cys*L), iron metabolism (*bes*A), antibiotic efflux (*mdt*A and *mdt*C), and β-lactam resistance (*bla*L1).

**Figure 4.**
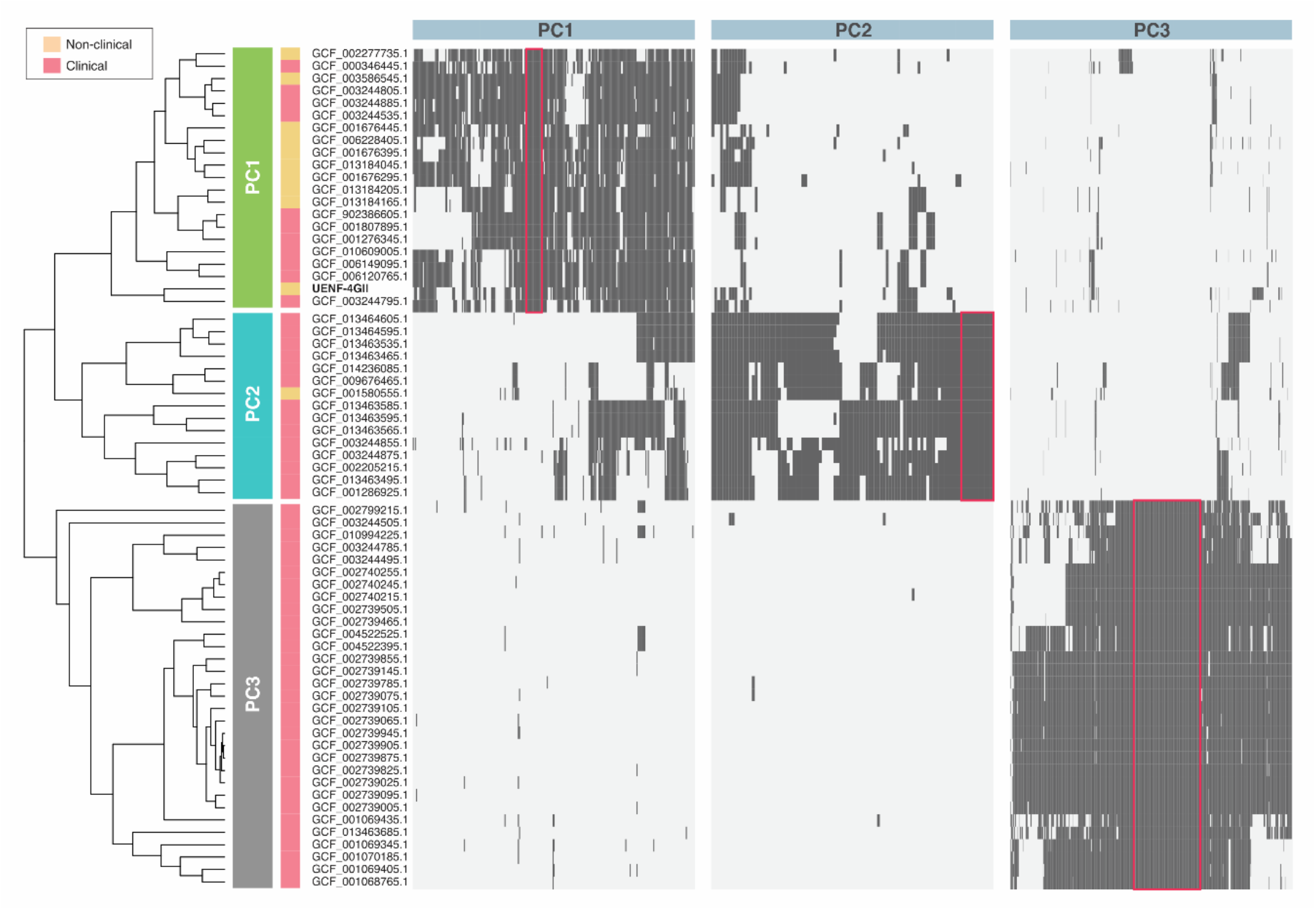
Distribution of PC-associated genes in *S. maltophilia* Sm3 genogroup. The cgMLST tree is annotated with two strips representing the phylogenetic cluster (PC) and the environment source. The heatmaps represent the presence (dark-gray) or absence (light-gray) of the genes identified by the pan-GWAS pipeline using phylogenetic cluster (PC) as trait. The neon red square highlights the most strongly associated genes found for each trait (i.e. 100% specificity and sensitivity).

The variants found in different traits (e.g. *bes*A, *mac*A and *bla*L1) are directly associated with genomic diversity of the PCs. A closer look into the *bla*L1 variant of PC3 reveals a molecular heterogeneity with 86% mean similarity with *bla*L1 from PC1 and PC2 genomes. Although the functional implications of such diversity is not fully understood, variation in *bla*L1 have been associated with β-lactam resistance in *S. maltophilia*, which may contribute to its increased prevalence as a nosocomial pathogen [63, 64, 69].

### Virulome and resistome analysis of the Sm3 genogroup

In order to understand the pathogenic potential within the Sm3 genogroup, we systematically investigated the distribution of virulence and antimicrobial resistance genes. The Sm3 virulome contains 54 genes (Supplementary table S6), out of which 35 constitute the core virulome. Most (21 out of 35) of the core virulence genes are involved in motility and adherence, including *pil*MNOPQ and the *pil*TU operon, which encode type IV pili subunits [70], and *tuf* (elongation factor Tu), involved in adhesion to host cells and extracellular matrix components [71]. In addition, we also identified seven type II secretion system (T2SS) genes (e.g. *xps*EFG and *hxc*RS), which promote the export of enzymes during bacterial colonization. Further, the presence of T2SS genes along with other core virulence genes (e.g. *acp*Xl, *hem*B, *hem*L, *csr*A, and *icl*), can be involved in infection and immune evasion capacity of Sm3 strains.

The Sm3 accessory virulome comprises 12 low and 7 high frequency genes, respectively (Figure 5). The high frequency genes are associated with adherence (*pin*N), motility (*flm*H), iron uptake (*bau*A), immune evasion (*rml*B), and stress tolerance (*clp*B, *kat*A, and *kat*G). The *clp*B gene encodes a heat shock-inducible chaperone required for bacterial tolerance to a variety of stresses, including heat, osmotic and acidic stress [72]. The *kat*A gene encodes a catalase that might confer tolerance against hydrogen peroxide-based disinfectants [18, 73] and is absent in all PC2 strains. Nevertheless, PC2 has an alternative catalase-peroxidase gene (*kat*G) that is absent in PC1 and PC3. The *kat*G gene is associated with oxidative stress control and has different roles in pathogenic bacteria [68, 74]. Four catalases (KatA1, KatA2, KatMn, and KatE) have been described in *S. maltophilia*, out of which KatA has been considered the main factor in conferring tolerance against hydrogen peroxide [75]. The apparent displacement of *kat*A by *kat*G in PC2 warrants further investigation and might provide insights the response of *S. maltophilia* to oxidative stress.

**Figure 5.**
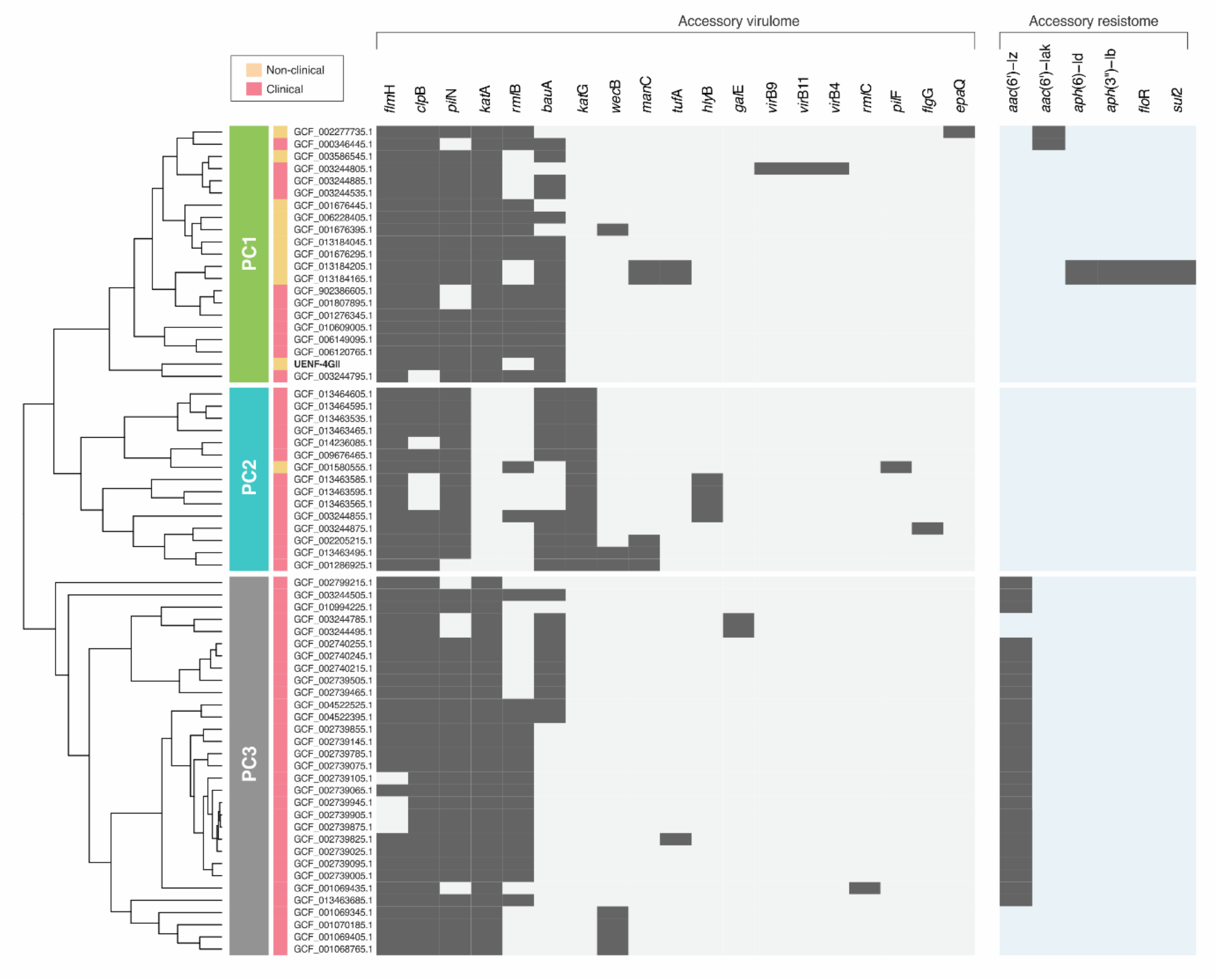
Acquired virulome and resistome of *S. maltophilia* Sm3 genogroup. The cgMLST tree is annotated with two strips representing the phylogenetic cluster (PC) identified and the environment source. The binary heatmaps represent the presence (dark-gray) or absence (light-gray/light-blue) of the genes identified.

The resistome analysis revealed 23 antimicrobial resistance genes (Supplementary table S7), out of which 17 comprise the core resistome, including the metallo-β-lactamase BlaL1 and the inducible Ambler class A β-lactamase BlaL2, as described above [76]. Further, antibiotic efflux pumps (e.g. SmeABC, SmeDEF, SmeRS, OqxAB, GolS, EmrE, and MexK) represent the main resistance mechanism encoded by the Sm3 core resistome.

The accessory resistome encompasses six genes (Figure 5). No accessory genes were found in PC2, as well as in most PC1 strains, including *S. maltophilia* UENF-4GII. The accessory resistome contains two classes of aminoglycoside-modifying enzymes: aminoglycoside phosphotransferase (encoded by *aph*(6)-Id and *aph*(3”)Ib); and N-aminoglycoside acetyltransferase (encoded by *aac*(6’)-Iz and *aac*(6’)-Iak). The *aac*(6’)-Iz gene was exclusively found in 25 (81%) PC3 genomes, whereas *aac*(6’)-Iak and *aph* genes were found in four PC1 genomes. Only two strains (G4S2 and G4S2-1) have the sulfonamide resistance gene *sul*2 and the chloramphenicol resistance gene *flo*R. In summary, the Sm3 genomes do not present a large repertoire of acquired resistance genes. The low frequency of sulfonamide resistance genes suggests that this antibiotic class might be a good therapeutic strategy against *S. maltophilia* infections, as previously proposed [18].

## CONCLUSION

In this study, we report the sequencing and in-depth analysis of the *S. maltophilia* UENF-4GII genome. This genome harbors a range of exclusive GIs associated with DNA-modification and anti-phage defense systems. Comparative analysis using pairwise genome identity metrics and cgMLST phylogeny show that *S. maltophilia* UENF-4GII belong to the Sm3 genogroup, which comprises three PCs. Using a pan-GWAS approach, we identified 131 genes as significantly associated with specific Sm3 PCs. Further, 471 genes were specifically associated with clinical and non-clinical environments. These genes could be used as biomarkers in future studies. Sm3 genomes comprise a low number of acquired virulence and resistance genes, although the presence of *kat*G and aminoglycoside resistance genes were associated with specific PCs. Collectively, our results provide important information regarding *S. maltophilia* genomic diversity that could provide the grounds for more detailed clinical and ecological investigations.

## ACKNOWLEDGEMENTS

This work was supported by Fundação Carlos Chagas Filho de Amparo à Pesquisa do Estado do Rio de Janeiro (FAPERJ; grants E-26/203.309/2016 and E-26/203.014/2018), Coordenação de Aperfeiçoamento de Pessoal de Nível Superior - Brasil (CAPES; Finance Code 001), and Conselho Nacional de Desenvolvimento Científico e Tecnológico. The funding agencies had no role in the design of the study and collection, analysis, and interpretation of data and in writing.

## AUTHOR CONTRIBUTIONS

Conceived the study: FP-S, TMV, FLO; Funding and resources: TMV, FLO; Data analysis: FP-S, HP-A; Interpretation of the results: FP-S, FPM, HP-A, TMV; Wrote the manuscript: FP-S, FPM, HP-A, TMV.

