## Supplementary figures for "Genome sequencing of the vermicompost strain *Stenotrophomonas maltophilia* UENF-4GII and population structure analysis of the *S. maltophilia* Sm3 genogroup"

[a] Laboratório de Química e Função de Proteínas e Peptídeos, Centro de Biociências e Biotecnologia, Universidade Estadual do Norte Fluminense Darcy Ribeiro (UENF), Brazil; [b] Departamento de Bioquímica e Imunologia, Instituto de Ciências Biológicas, Universidade Federal de Minas Gerais, Belo Horizonte, MG, Brazil; [c] Núcleo de Desenvolvimento de Insumos Biológicos para a Agricultura (NUDIBA), UENF, Brazil; [d] Laboratório de Biologia Celular e Tecidual, Centro de Biociências e Biotecnologia, UENF, Brazil.

*** Corresponding author:**

Thiago M. Venancio; Laboratório de Química e Função de Proteínas e Peptídeos, Centro de Biociências e Biotecnologia, Universidade Estadual do Norte Fluminense Darcy Ribeiro (UENF); Av. Alberto Lamego 2000, P5 / sala 217; Campos dos Goytacazes, Rio de Janeiro, Brazil..

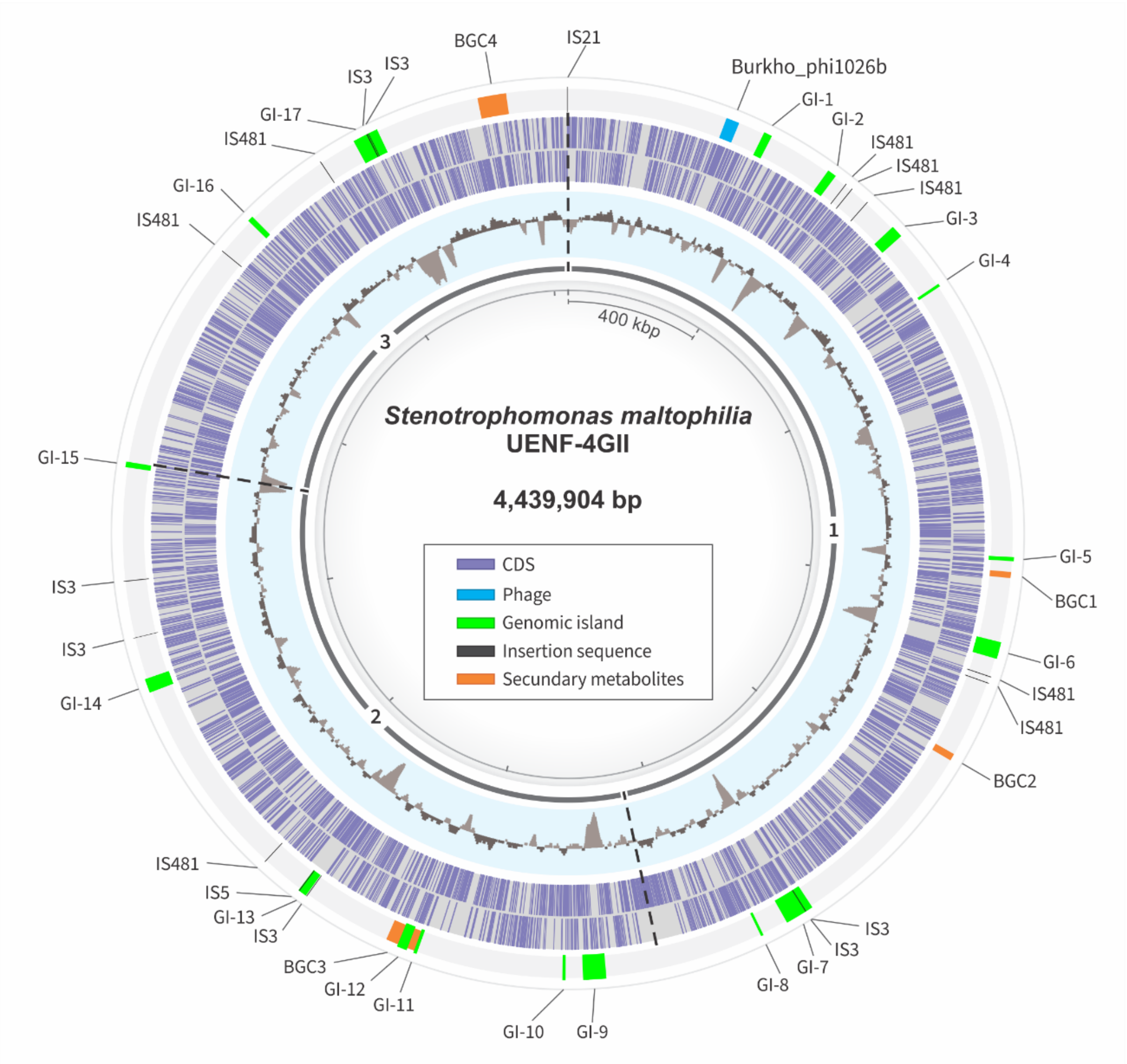


**Supplementary figure S1. Circular representation of the *S. maltophilia* UENF-4GII genome.** From inner to outer ring: the first ring represents the scaffold number. The second ring represents GC content; light-grey indicates GC content higher than average; dark-grey indicates less than average. The third circle ring represents predicted CDS on the plus and minus strands, respectively. The fourth ring represents genome features such as phage, genomic islands, insertion sequence and secondary metabolites.

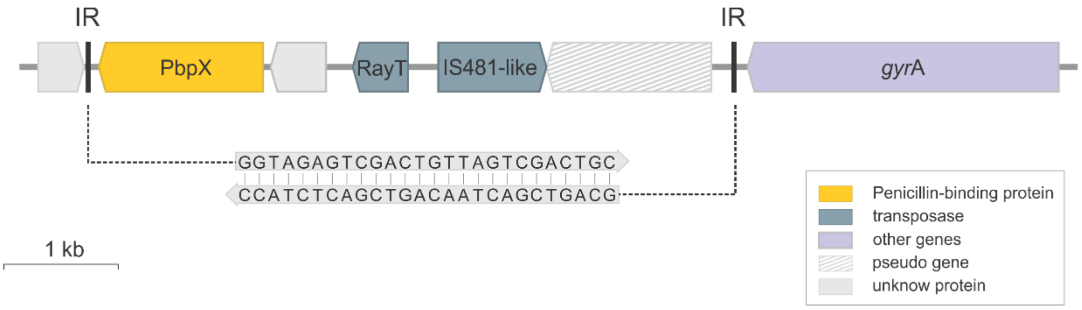


**Supplementary figure S2. Genetic context of *S. maltophilia* UENF-4GII composite transposon.** The IR represents two perfect 26-bp inverted repeat sequence. The transposon contains a *pbp*X gene winch encodes a penicillin-binding protein associated with penicillin resistance; two different transposases-encoding genes; and two unknown function genes.

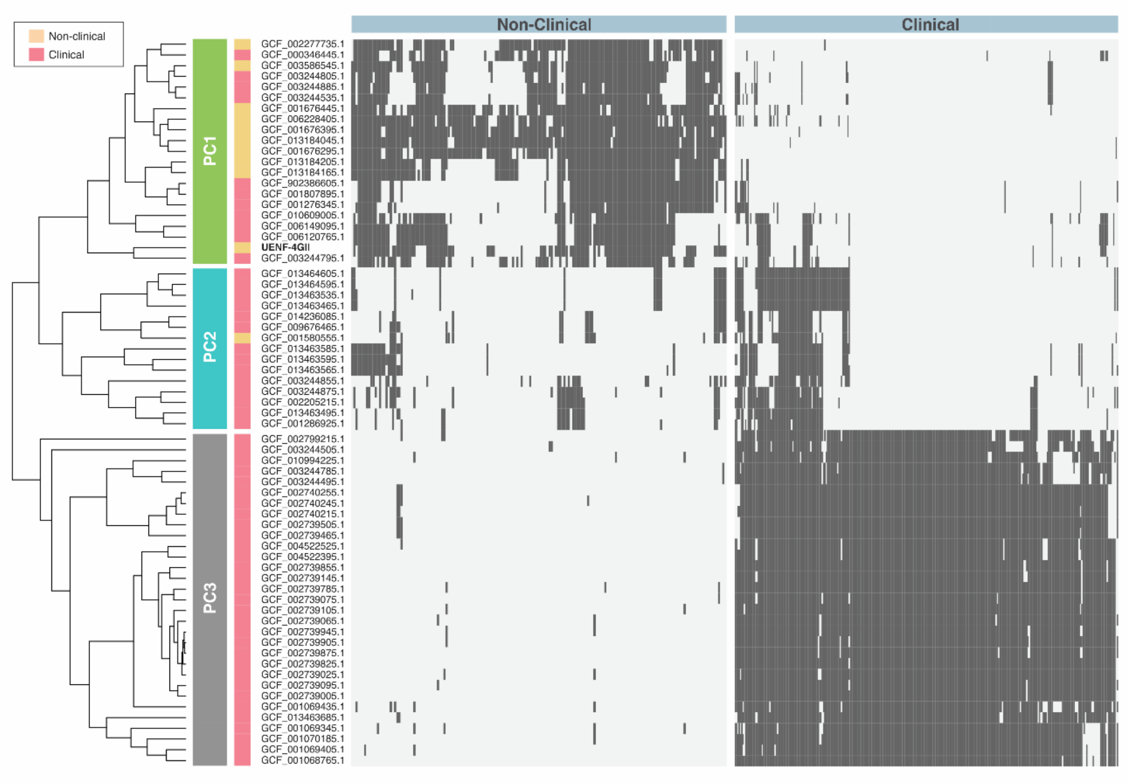


**Supplementary figure S3. Distribution of environmental-associated genes in Sm3 genogroup.** The cgMLST tree is annotated with two colored strips representing the phylogenetic cluster (PC) and the environment source. The heatmaps represent the presence (dark-gray) or absence (light-gray) of the genes identified by the pan-GWAS pipeline using the environment source as trait.

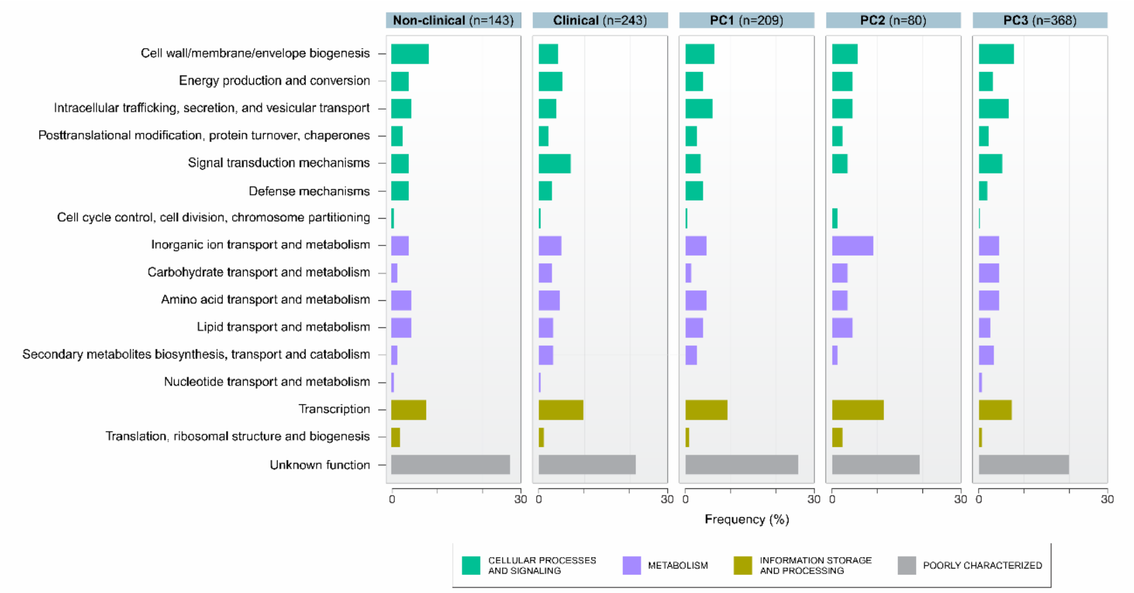


**Supplementary figure S4.** Functional categories identified by EggNOG annotation of environment and PC-associated genes.
